# GNOM-Dependent Endosomal Recycling Sets the Timing of Founder Cell Specification

**DOI:** 10.1101/2023.01.31.526479

**Authors:** Pablo Perez-Garcia, Juan Perianez-Rodriguez, Alvaro Sanchez-Corrionero, Miguel A. Moreno-Risueno

## Abstract

Lateral root growth occurs from primed xylem pole pericycle cells that can be specified as founder cells, but how these cells acquire the pluripotency state to initiate a root organogenesis program is not fully understood. Here, we find that the EMS 204 mutant is unable to specify founder cells due to a point mutation affecting the C-terminal HDS3 domain of the GNOM protein. We used genetics and cell biology to study the trafficking machinery of xylem pole pericycle cells, cells in which GNOM is primarily associated with recycling endosomes. We found that the lack of the HDS3 domain negatively affects the stability of the GNOM protein within the cells of the xylem pole pericycle and, subsequently, the integrity of the trafficking machinery of these cells. We determined that GNOM is responsible for the recycling of AUX1 toward the plasma membrane at the xylem pole pericycle and impairment of this recycling results in premature accumulation of AUX1 in the vacuole. In this work, we describe a cell type-specific cellular mechanism and the role of a particular domain during lateral root development.

## 2. INTRODUCTION

The timing of founder cell (FC) appearance along the root is based on a mechanism known as the Root Clock (RC) (1). RC periodicity results in oscillating peaks of auxin maximum at the root tip delineating an area known as the oscillation zone (OZ) (1). RC entrainment depends on a bistable auxin module, although auxin fluxes derived from the lateral root cap also regulate the pace of the clock (2-4). Not only auxin but other metabolites such as retinoids modulate the rhythmicity of the RC or the accumulation of esterified pectin in a GNOM-dependent manner (5, 6). The oscillatory behavior of the RC determines the position of pre-branch sites (PBS) along the primary root that periodically prime xylem pole pericycle (XPP) cells with competence to be specified as FC later (1, 2, 5, 6).

Most of the candidate factors to regulate the cellular conversion of XPP cells to pluripotent FC described to date are regulated by the hormone auxin, although the function of these regulators seems to be immediately after the acquisition of FC identity (7-10). Exceptionally, MEMBRANE ASSOCIATED KINASE REGULATOR 4, whose expression also depends on auxin levels, is expressed in XPP before specification to FC and alters the root architecture so it could play a role during this process (2). After the FCs are specified, coordinated nuclear migration occurs within two adjacent FCs, indicating dramatic cellular reorganization (7). Interestingly, the described regulators of this process also share upstream auxin regulation, suggesting that an auxin stimulus may be necessary during FC specification just before nuclear migration (7-9). Consistently, microscopic studies to monitor auxin signaling also suggest a second auxin peak within XXP-primed cells before specifying as pluripotent FC (11), although the cellular mechanism remains unresolved.

FCs can divide asymmetrically to produce daughter cells with distinct morphology and function, indicating their pluripotent properties (12). Pluripotency and auxin signaling are commonly linked (13-17), for example, when pericycle-like cells distributed throughout the plant act as pluripotent cells upon application of auxin (16). Although evidence supports a close relationship between pluripotency acquisition and auxin signaling, the mechanism that determines FC pluripotency acquisition is unknown.

The GNOM protein is a large guanine nucleotide exchange factor (GEF) that acts on ADP-ribosylation factor (ADP), small GTPases involved in vesicle trafficking (18-21). GEF-ARFs associate with endomembranes when they bind to GTP, triggering the recruitment of coat proteins for vesicle establishment (22). GNOM is found primarily in the Golgi apparatus, although it maintains the Trans Golgi Network (TGN) structure that affects vesicular recycling to the plasma membrane (23). Structurally, in the N-terminal half of the GNOM protein, the cyclophilin-binding domain (DCB) is responsible for homodimerization and the sec7 domain for biochemical function (21, 24, 25). The C-terminal part is composed of three HDS domains (homology downstream of Sec7) (26), but the deletion of this C-terminal half of the GNOM protein does not show phenotypes as dramatic as those found in mutants affecting the N-terminal portion of the protein (27). Until date, no specific function has been attributed to HDS domains.

GNOM mediates asymmetric trafficking of the auxin efflux transporter PIN1 to the basal plasma membrane of the cell (19, 20). This is essential to establish the auxin gradient for proper acquisition of cell identities during embryogenesis, therefore several *GNOM* mutants result in embryonic lethality (18, 19, 28). GNOM is also involved in auxin transport during post-embryonic development, especially in primary root growth and lateral root formation (27, 29-32). PIN1 is expressed from the earliest stages of primordium formation, suggesting a role for GNOM in orchestrating the accumulation of PIN1 in its functional location at this stage (31, 32). Inducible repression of GNOM translation also showed pericycle-related postembryonic effects of GNOM (32). In agreement, GNOM alleles affected at their C-terminus have difficulty reaching the necessary auxin levels at the pericycle to develop a lateral root primordium (LRP) (31). Although the maximum amount of auxin needed to initiate a lateral root dependents on GNOM, specifically its C-terminal half, GNOM-dependent transport of PIN1 has not been linked to this process.

Here, we characterize the EMS-generated mutant *204* and find its inability to specify FC. Next generation sequencing revealed a point mutation in the HDS3 domain of GNOM as the mutation responsible for the observed phenotype. We found a cell-specific disruption of the trafficking machinery in XPP cells. Consistently, we observed the GNOM protein primarily associated with recycling endosomes within XPP cells and in FCs. A recombinant GNOM protein that mimicked the EMS-derived version of GNOM produced in the mutant *204* showed decreased GNOM stability in XPP and FC cells. We found that the trafficking of the auxin influx transporter AUX1, previously thought to be GNOM-independent, was severely impaired in the *204* mutant. Furthermore, we showed that AUX1 accumulated prematurely in vacuoles in the *204*-mutant XPP cells. In this article, we describe how specific trafficking of the auxin transporter AUX1 to the plasma membrane in XXP cells mediated by the HDS3 domain of GNOM is a necessary mechanism by which FC specifies.

## 3. RESULTS

### The GNOM HDS3 domain is required during FC specification

To search for affected plants during founder cell (FC) specification, we reasoned that these plants should be completely unable to proceed to the subsequent steps to develop lateral roots (LR). An EMS mutagenesis screen allowed us to find the *204* mutant that lacked LR and displayed compromised root length (Fig. 1a, 1c–d). Next-generation sequencing revealed a single-nucleotide polymorphism (SNP) in the C-terminal domain HDS3 of GNOM, a protein known to be involved in cellular trafficking (20, 29, 33, 34). The point mutation changed a cytosine at position 3492 to a thymine, causing a premature *STOP* codon and resulting in almost complete deletion of the HDS3 domain (Fig. 1b). To determine whether this mutation was responsible for the observed phenotypes, we introduced the genomic portion of *GNOM* into the *204* mutant. Quantification of root length and lateral root formation showed recovery by complementation of the genomic *GNOM* into the *204* mutant. (Fig. 1a, 1c-d). These results indicate that GNOM is indeed the mutated gene responsible for the phenotypes described in the *204* mutant. To test the effect of the *204* mutation on FC specification, we introduced the FC cell specification marker *pSKYP2B:NLS-3xmCherry* (35) that is expressed in cells of the xylem pole pericycle (XPP) before nuclear migration. We were unable to detect *pSKYP2B:NLS-3xmCherry* expression in mutant *204*, but *pSKP2B:NLS-3xmCherry* expression was recovered in complemented plants, indicating that this mutant is impaired in FC specification (Fig. e)

**Fig1.**
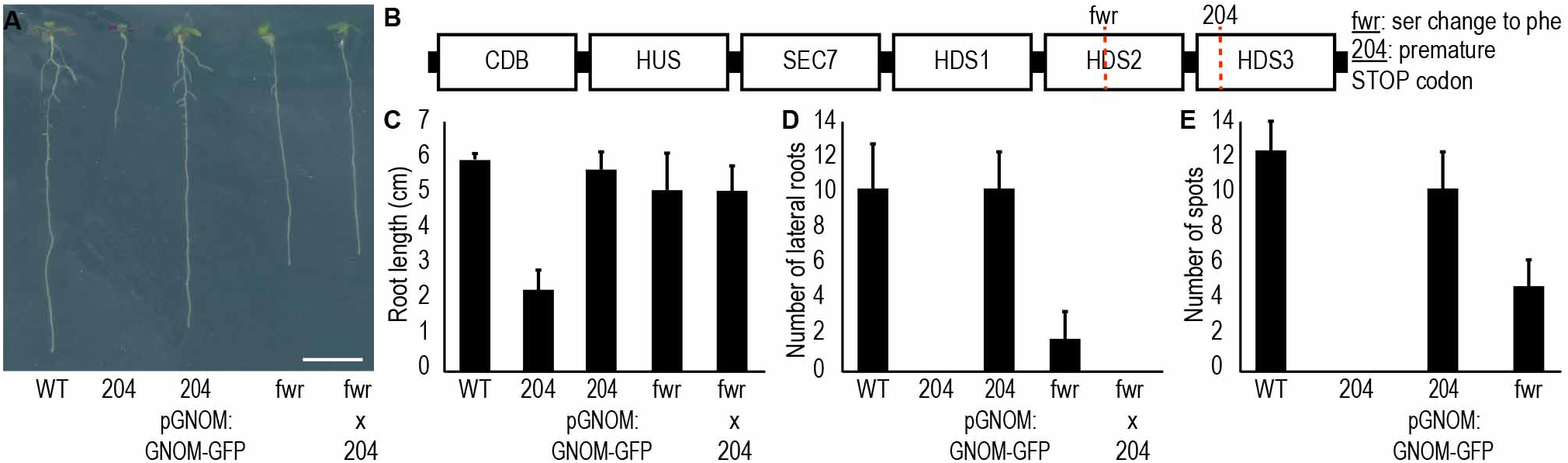
The *204* mutation on *GNOM* gene prevents founder cell specification. **(a)** Picture of WT, *204* mutant, *204* mutant complemented with *pGNOM:GNOM-GFP* construct, *fewer* (*fwr*) mutant and trans-heterozygous *204/fwr* mutant seedlings at 8 days post inhibition (dpi). **(b)** Schematic representation of the GNOM protein indicating the relative position of the GNOM domains and the location of the *204* and *fwr* mutation. (c,d,e) Graphs showing root length at 8 dpi **(c)**, lateral roots quantity **(d)** and number of *pKYP2B::ER/NLS-3x-Cherry* in the primary root **(e)**.

To further determine the specific role of the HDS3 domain, we compared available *GNOM* mutants in similar regions of the coding sequence. Another *GNOM* allele with a single amino acid change near the HDS3 domain was characterized and named *fewer* (*fwr*) based on its reduced number of lateral roots (Fig. 1a–b) (31). Introgression of *pSKYP2B:NLS-3xmCherry* into *fwr* mutant plants confirmed the functional, albeit severely reduced, ability of this mutant to specify FC, in contrast to the *204* mutant (Fig. 1e). We used *204/fwr* transheterozygotes to determine the strength relationship between these alleles, which could help us better understand the actual relevance of the HDS3 domain during FC specification. Root length was similar in homozygous *fwr* than in trans-heterozygous *204/fwr*, although the few LRs developed in the *fwr* mutant were lost in the trans-heterozygous 204/fwr mutants, resembling the *204* phenotype (Fig. 1a–d). These findings evidence the stronger effect of the *204* mutation during LR formation. Complete deletion of the HDS3 domain prevents FC specification, whereas amino acid point mutations near the HDS3 domain only reduce the number of specified FCs, but do not completely prevent their appearance. Taken together, these results highlight the importance of a complete and uncompromised HDS3 domain of the GNOM protein in maintaining its functionality during FC specification.

### The XPP cell trafficking machinery is compromised in the *204* mutant

GNOMs have been studied extensively in recent years due to their biological importance in regulating the integrity of the trafficking machinery and vesicular trafficking during development (19, 20, 27, 28, 31, 33, 34). As FC specifies from XPP-primed cells, we hypothesized that if GNOM-dependent trafficking was impaired in XPP cells in the *204* mutant, we should observe structural changes within these cells in the endomembranes where GNOM usually associates (20, 23). In fact, we could appreciate changes in the pattern of the Golgi apparatus in the *204* mutant in XPP cells (Fig. 2a-b). When comparing the pattern of the marker *pUBQ10::RFP-Rab A5d* (36), which is associated with recycling endosomes, in WT and *204* XPP mutant cells, the differences became even more clear (Fig. 2c–d). These results indicate that the lack of the HDS3 domain in the *204* mutant has a predominant effect on the organization of the XPP cells endomembrane system.

**Fig2.**
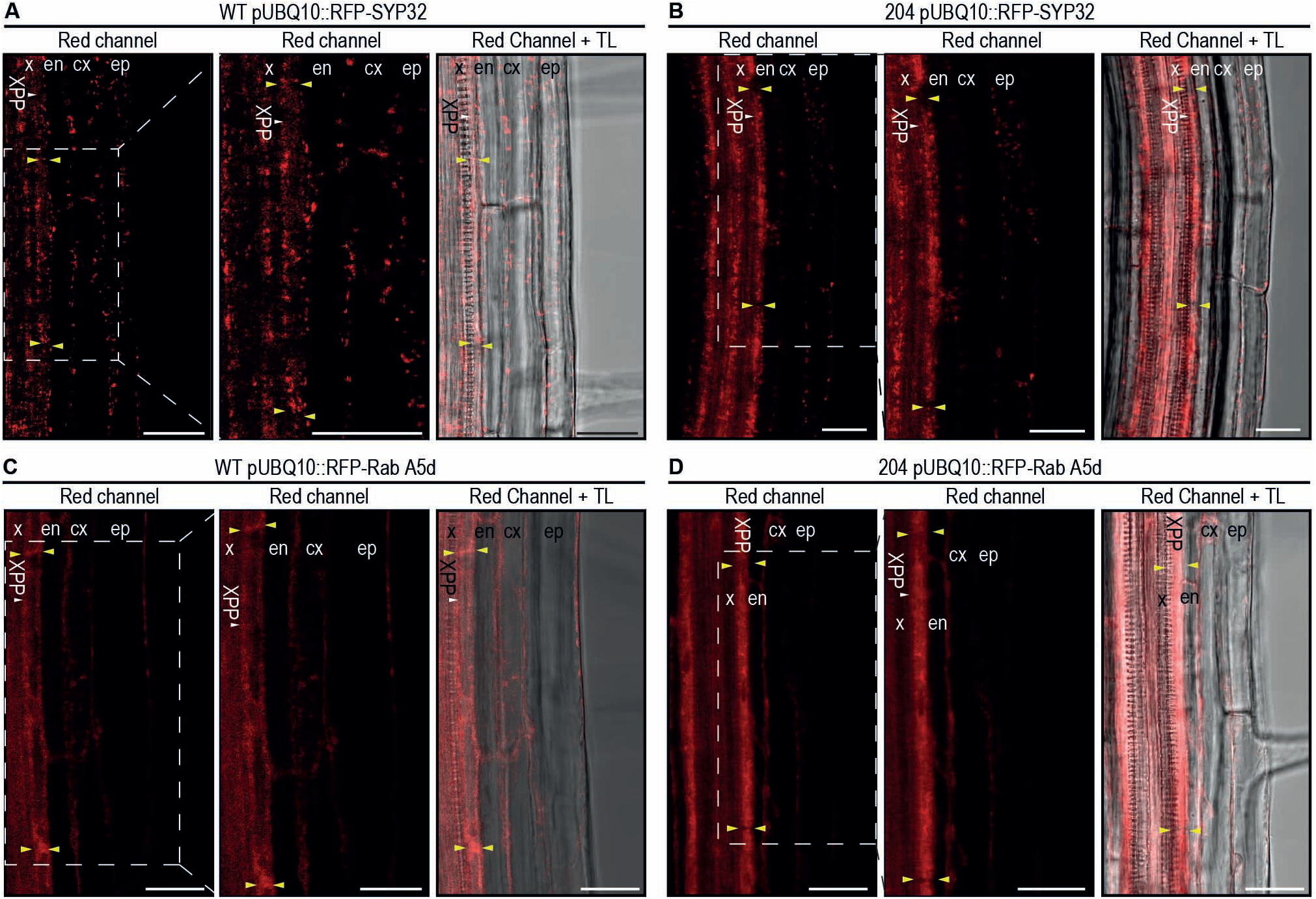
Trafficking machinery is disorganized in cells of the xylem pole pericycle in the 204 mutant. **(a, b)** Confocal images showing the Golgi apparatus marker *pUBQ10::RFP-SYP32* in roots with focus on the xylem pole pericycle (XPP) cells in WT (a) and *204* mutant plants (b). **(c, d)** Confocal images showing the traffic endosome marker *pUBQ10::RFP-Rab A5d* in roots with focus on the XPP cells in WT (c) and *204* mutant plants (d). White arrowheads: XPP cells. Yellow arrowheads: apexes of the same XPP cell. ep: epidermis. cx: cortex. en: endodermis. x: xylem. TL: transmitted light. Scale bars: 25 μm.

### GNOM primary associates with recycled endosomes in XPP cells

Then we studied the dynamics of GNOM during the specification of the FC. As GNOM has been linked to the integrity of the trafficking machinery (20, 23, 37, 38) and we observed disrupted patterns of the Golgi apparatus and recycling vesicles in XPP cells of the *204* mutant (Fig. 2a–d), we decided to investigate whether GNOM was associated with these endomembranes in XPP cells. We used transgenic plants carrying the genomic version of *GNOM* fused to a *GFP* and under the regulation of its own promoter (*pGNOM::GNOM-GFP*). GNOM has been observed to associate primarily with Golgi membranes (23). In agreement, we found high colocalization of Golgi aggregations with GNOM-GFP protein in epidermal cells of the meristem, but in XPP cells this colocalization was severely reduced (Fig. 3a–b). In contrast, we observed increased colocalization of GNOM-GFP protein with recycling endosomes in XPP cells (Fig. 3c), which could explain the dramatic disorganized pattern of these organelles observed in these cells in the *204* mutant (Fig. 2c–d). These findings most likely indicate a major role of GNOM during the recycling of cargo toward the membrane in cells of the XPP before their specification to FC.

**Fig3.**
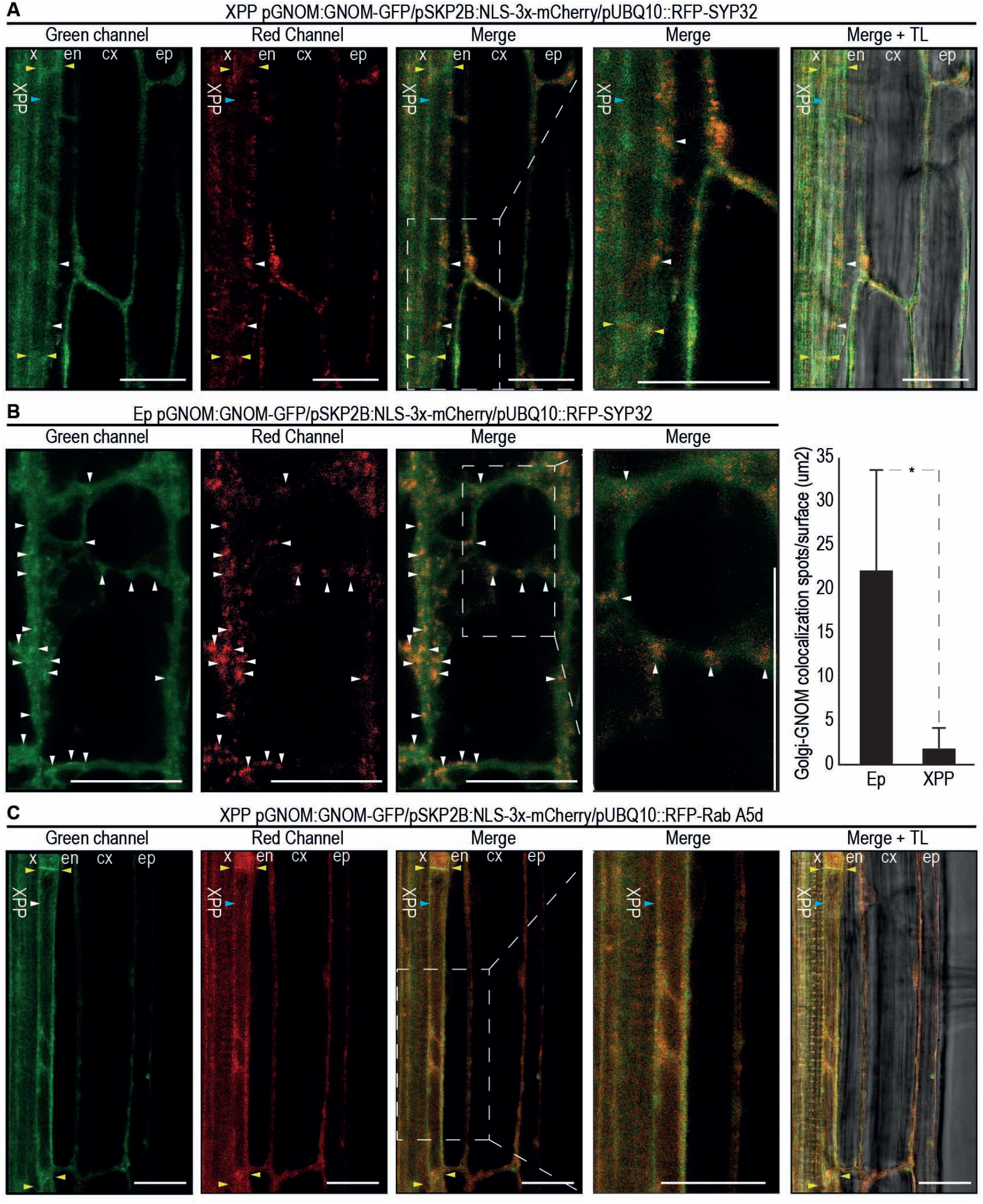
GNOM primary associates with recycling endosomes in the XPP. **(a)** Confocal images showing the accumulation of the GNOM-GFP recombinant protein and the Golgi apparatus marker *pUBQ10::RFP-SYP32* in roots with focus on the a XPP cell in WT. **(b)** Confocal images showing the accumulation of the GNOM-GFP recombinant protein and the Golgi apparatus marker *pUBQ10::RFP-SYP32* in epidermal cells of the meristem with focus on an specific area of an epidermal cell in WT plants. Right panel: graph comparing the number of Golgi accumulates found in the same area in XPP cells and epidermal cells in WT plants. **(c)** Confocal images showing the accumulation of the GNOM-GFP recombinant protein and the traffic endosome marker *pUBQ10::RFP-Rab A5d* in roots with focus on a XPP cell in WT. Yellow arrowheads: apexes of the same XPP cell. White arrowheads: Golgi accumulates. ep: epidermis. cx: cortex. en: endodermis. x: xylem. TL: transmitted light. Scale bars: 25 μm. (b) n≥ 10. Error bars: S.D. Asterisk: p-value < 0.001 by General Linear Model

To verify this idea, we monitored the accumulation of GNOM in FC and after the first asymmetric division that occurs during the development of lateral root primordia (LRP) (7-9). We first observed Golgi aggregates in parallel with the GNOM-GFP protein after the specification of two adjacent FCs, tagged with the *pSKYB2B:NLS-3xmCherry* marker, and after the first asymmetric division. As expected from our previous observations, there was little co-location of GNOM-GFP protein and Golgi aggregates at these developmental stages (Fig. 4a–b). When we monitored the dynamics of recycling endosomes together with GNOM-GFP protein accumulation, we observed high spatiotemporal colocalization during FC specification and after the first asymmetric division (Fig. 4c–d). Taken together, these results demonstrate the major role of GNOM recycling proteins toward the plasma membrane before and during FC specification.

**Fig4.**
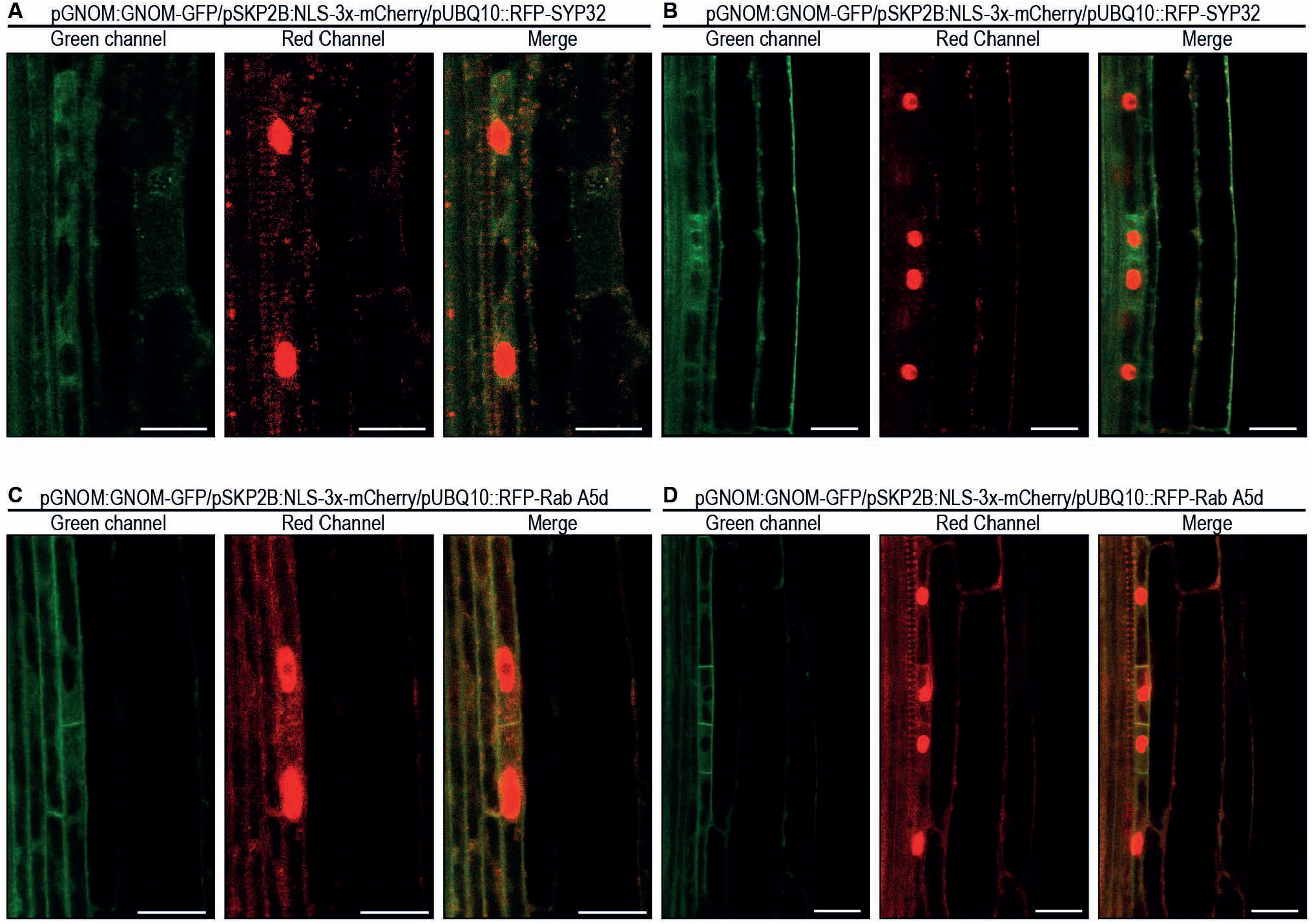
GNOM primary associates with recycling endosomes after the specification of FC. **(a, b)** Confocal images showing the accumulation of the GNOM-GFP recombinant protein and the Golgi apparatus marker *pUBQ10::RFP-SYP32* in two adjacent FC (a) and after the first asymmetric division (b). **(c, d)** Confocal images showing the accumulation of the GNOM-GFP recombinant protein and the traffic endosome marker *pUBQ10::RFP-Rab A5d* in two adjacent FC (c) and after the first asymmetric division (d).

### The HDS3 domain stabilizes GNOM in XPP cells

Although the GNOM protein family is very conserved among kingdoms, the HDS domains are not found in other eukaryotic proteins and their role is unclear (26). We wondered if the lack of the HDS3 domain might be interfering with the subcellular localization of GNOM and its function in the XPP cells. To address this question, we generated a version of the GNOM protein fused with a YFP after the site of the premature STOP codon generated by the SNP mutation responsible for the *204* phenotype and regulated under its own promoter (*pGNOM::GNOM204version-YFP*). We first transiently introduced this construct and the full version of GNOM (*pGNOM::GNOM-GFP*) into tobacco leaves. We observed the accumulation of both versions in the Golgi endomembranes (Fig. S1a–b), suggesting that the presence of these proteins in another compartment in *Arabidopsis* may involve other partners and further regulation. Of note, the *pGNOM::HDS3-YFP* construct introduced into tobacco leaves was also found Golgi-associated in agreement with GNOM showing strong association with this organelle in the absence of an additional cellular environment (Fig. S1c).

When we specifically observed XPP cells in *Arabidopsis*, we did not detect GNOM204version-YFP protein, which could explain the defects in the trafficking machinery in the *204* mutant predominantly affecting XPP cells. Even after the first asymmetric division, when GNOM-GFP protein accumulation became evident in the small daughter cells (Fig. 4b, 4d), we did not see any YFP signal of the GNOM204version-YFP protein (Fig. 5a–b). These observations suggested that the HDS3 domain might favor GNOM stability and thus, the full version of GNOM, including the HDS3 domain, is more stable in XPP cells.

**Fig5.**
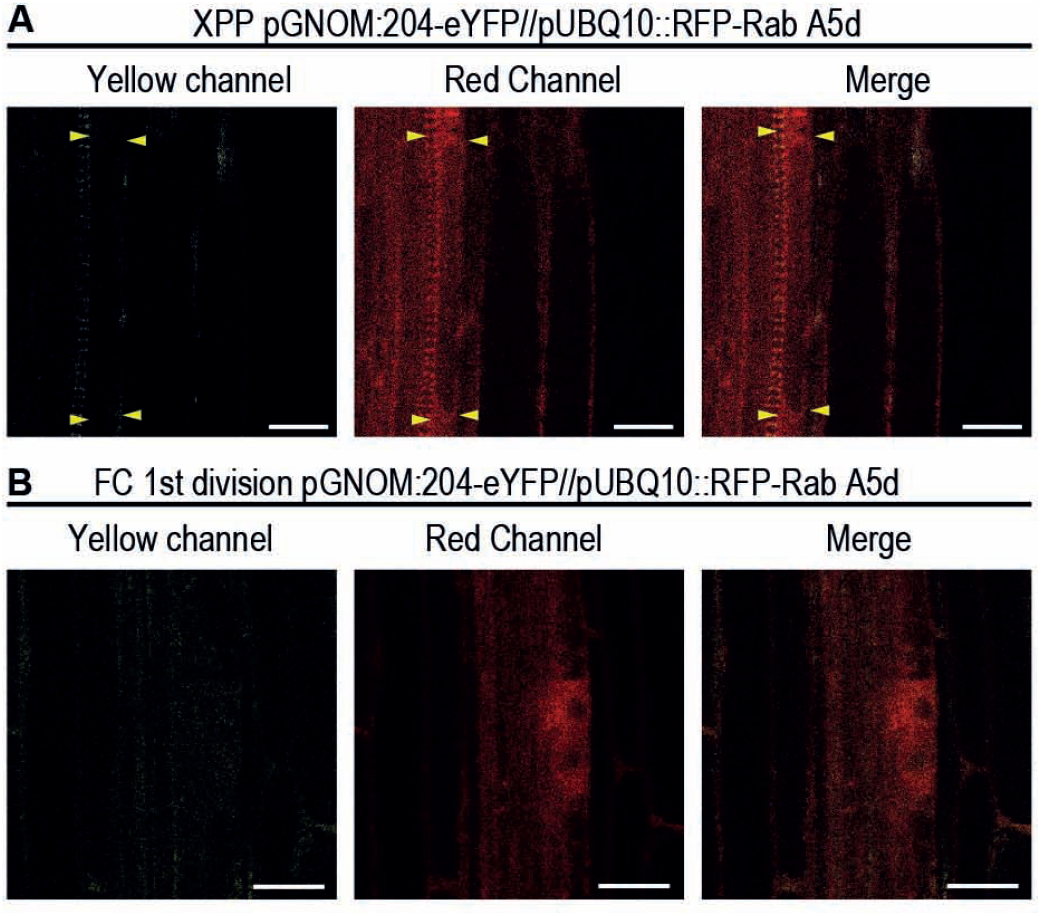
The HDS3 domain of GNOM contributes to GNOM protein stability in the XPP cells. **(a, b)** Confocal images showing the accumulation of the GNOM-204-eYFP recombinant protein and the traffic endosome marker *pUBQ10::RFP-Rab A5d* in a XPP cell (a) and after the first asymmetric division (b). Yellow arrowheads: apexes of the same XPP cell. Scale bars: 25 μm.

### Localization of AUX1 is disrupted in XPP cells in the *204* mutant

Phenotypes of the mutant *gnom* have been linked to disrupted auxin fluxes due to the inability of polar auxin transporters to reach their functional subcellular locations in the absence of a functional GNOM protein (19, 20, 27, 28, 31, 32). For this reason, we explored whether the root development phenotypes present in the *204* mutant could be related to problems in the trafficking of polar auxin transporters. We supplemented *204* mutant plants with auxin to try overcoming the lack of LR development, and indeed 204 auxin-supplemented mutant plants could produce LR (Fig. 6a). These results could indicate that the lack of the GNOM HDS3 domain causes a problem with auxin fluxes within XPP cells that prevent their specification to FC.

**Fig6.**
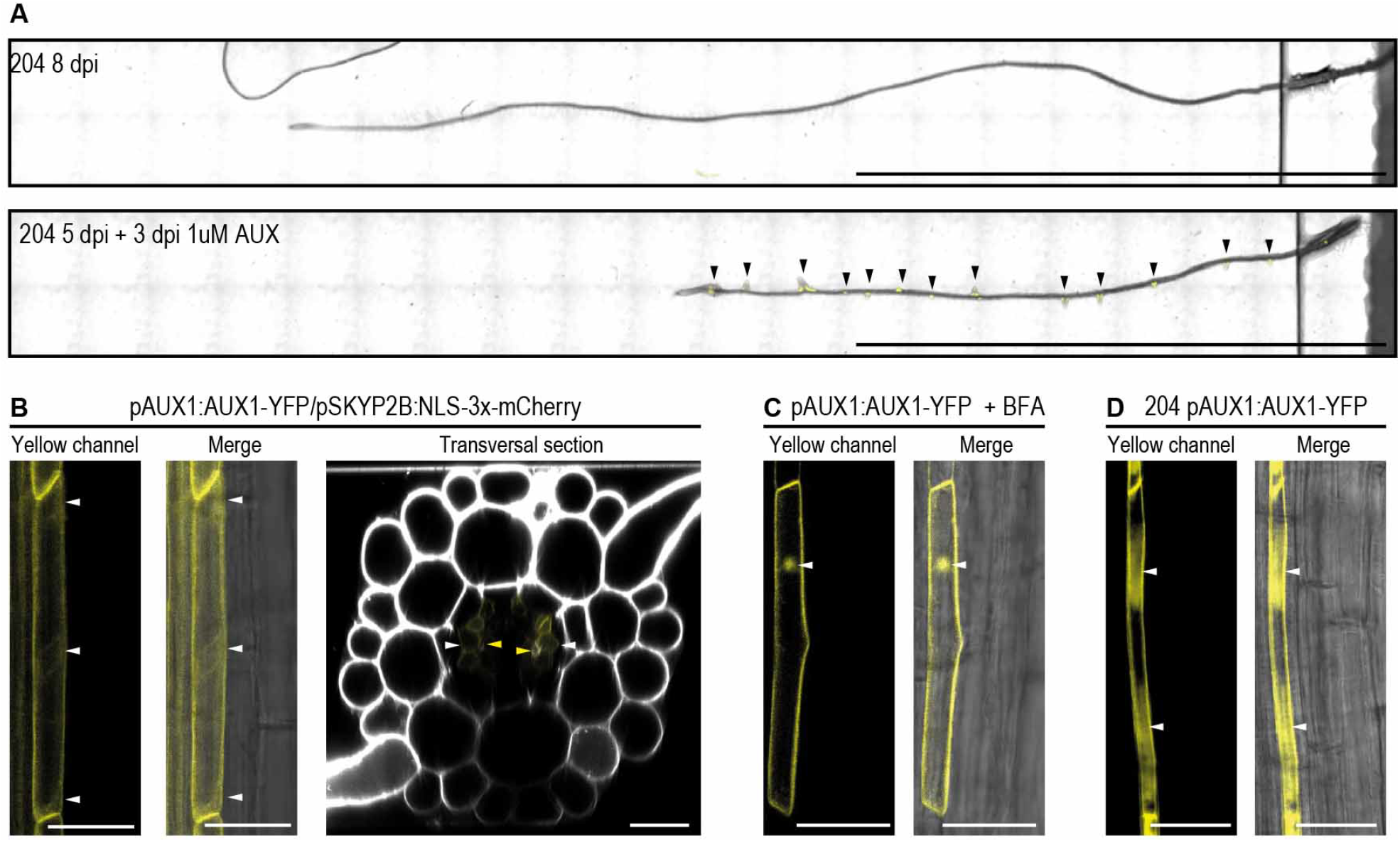
Subcellular localization of AUX1 and PIN3 is altered in the *204* mutant. **(a)** Confocal images of WT and *204* mutant roots with and without auxin supplementation. **(b, c, d)** Confocal images showing AUX1-YFP recombinant protein accumulation in a XXP cell and in a transversal section of WT roots (a), in a XPP cell after treatment with brefeldin A (BFA) (b) and in a XPP cell of the *204* mutant (c). Black arrowheads: Lateral roots. White arrowheads: sites of AUX1-YFP recombinant protein accumulation. Yellow arrowheads: Xylem. Scale bars: 1 cm (a) 25 μm (b, c, d)

GNOM is involved in the trafficking of the polar auxin transporter PIN1 (19, 20, 31, 39), therefore we decided to monitor PIN1 accumulation during FC specification. We detected PIN1-GFP protein at the division plane of two adjacent newly specified FCs (Fig. S2a–b), indicating that PIN1 trafficking could not explain the lack of specified FCs found in the *204* mutant.

We then studied the protein accumulation pattern of the auxin importer AUX1, although its membrane trafficking has been reported to be GNOM-independent (20, 40). We used *pAUX1:AUX1-YFP* transgenic lines and found AUX1-YFP associated with XPP cells from the elongation zone to the differentiation zone (Fig. 6b). To determine whether GNOM may be involved in the subcellular localization of AUX1 in XPP cells, we decided to treat plants with Brefeldin A (BFA), an inhibitor of GNOM and GNOM-like proteins (41). BFA treatments in *pAUX::AUX1-YFP* plants resulted in the accumulation of AUX1-ÝFP in so-called BFA bodies (20, 41) (Fig. 6c), suggesting that AUX1 trafficking is regulated by GNOM in the XPP. Consistently, introgression of *pAUX::AUX1-YFP* into the *204* mutant resulted in delocalization of AUX1-YFP in XPP cells (Fig. 6d). Interestingly, this effect was cell-type specific, as we did not detect any difference in AUX1-YFP localization in any meristematic cells under BFA treatment or in the *204* mutant background in concordance with previous reports (Fig. S3). Taken together, these findings demonstrate a cell-type role of GNOM in AUX1 trafficking that dependents on the C-terminal HDS3 domain of GNOM.

### Accumulation of AUX1 in the vacuole is accelerated in the 204 mutant

We have described GNOM-dependent trafficking of AUX1 within XPP cells, thus we hypothesize that AUX1 should be dynamically recycled in this cell type. To test this hypothesis, we crossed *pAUX1::AUX1-YFP* plants with the marker for recycling endosomes *pUBQ10::RFP-Rab A5d* and, similar to what we observed for the GNOM-GFP protein (Fig. 3c), AUX1-YFP protein is mostly found associated with these vesicles in XPP cells (Fig. 7a). Interestingly, AUX1-YFP remains excluded from vacuoles in this cell type (Fig. 7b), but during the early stages of LRP development, AUX1-YFP accumulates within vacuoles (Fig. 7c). This observation led us to the idea that it might be possible that, in the case of the *204* mutant, AUX1 could be delivered to the vacuole earlier in development, preventing AUX1 from being recycled to the XPP cell membrane. We first confirmed that AUX1-YFP accumulated in the vacuole of XPP cells in the *204* mutant by crossing *pAUX1::AUX1-YFP* plants with the vacuole marker *pUBQ10::RFP-VAMP711* (42) (Fig. 7d). To verify that this accelerated accumulation of AUX1 in the vacuole was due to deficiencies in GNOM-dependent recycling of AUX1 to the membrane, we treated *pAUX1::AUX1-YFP* plants with the recycling inhibitor endosidine2 (EN2) (43). We observed a *204* mutant-like accumulation of AUX1-YFP proteins in the vacuoles (Fig. 7e), confirming that the lack of recycling of AUX1 toward the plasma membrane is what causes its premature accumulation in this organelle.

**Fig7.**
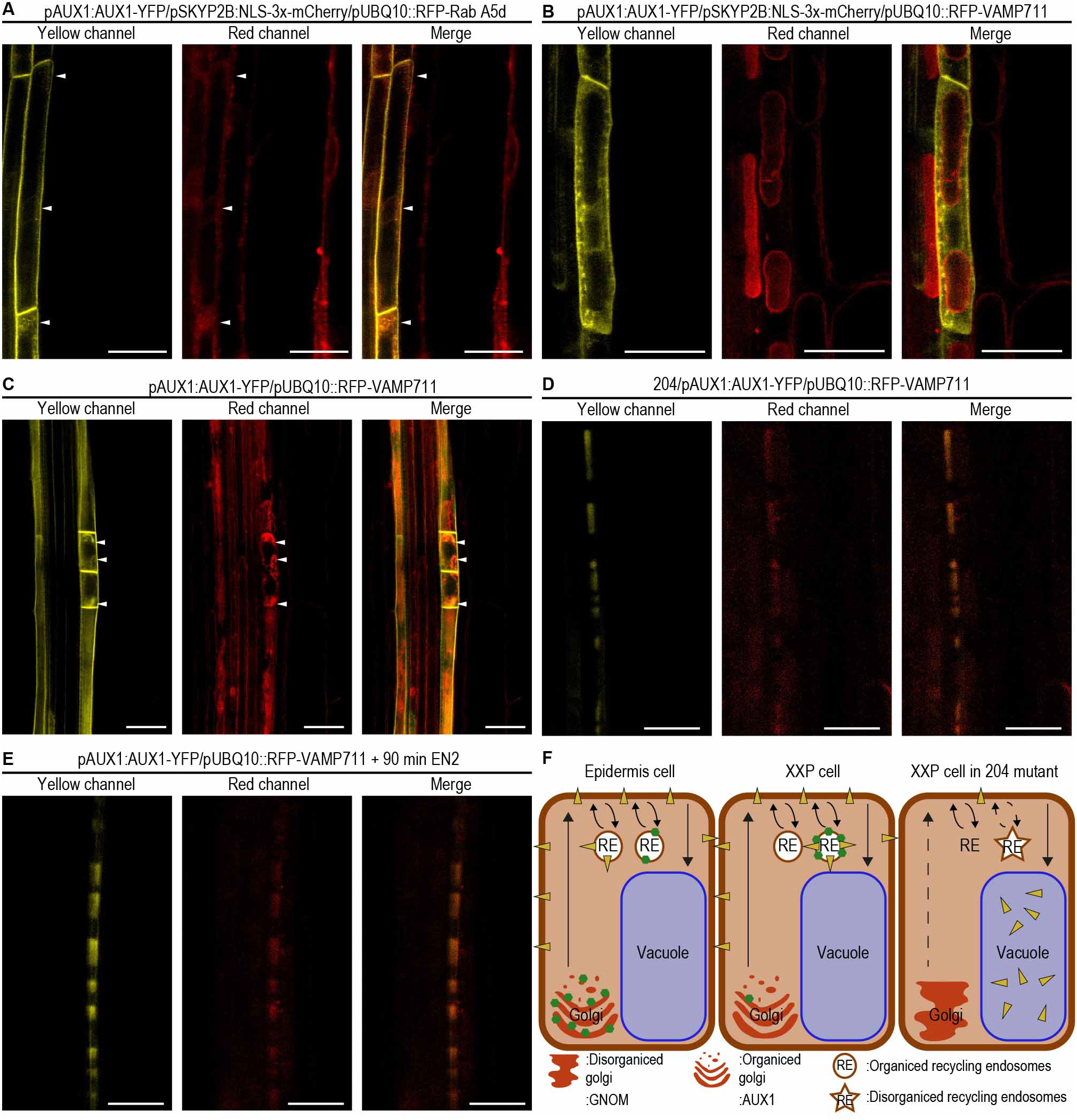
Decreased protein recycling in the 204 mutant causes premature accumulation of AUX1 into the vacuole. **(a)** Confocal images showing the accumulation of the AUX1-YFP recombinant protein and the traffic endosome marker *pUBQ10::RFP-Rab A5d* in a XPP cell in a WT plant **(b)** Confocal images showing the accumulation of the AUX1-YFP recombinant protein and the vacuole marker *pUBQ10::RFP-VAMP711* in a XPP cell in a WT plant. **(c)** Confocal images showing the accumulation of the AUX1-YFP recombinant protein and the traffic endosome marker *pUBQ10::RFP-Rab A5d* after the first asymmetric division in a WT plant. **(d)** Confocal images showing the accumulation of the AUX1-YFP recombinant protein and the vacuole marker *pUBQ10::RFP-VAMP711* in a XPP cell in a *204* mutant plant. **(e)** Confocal images showing the accumulation of the AUX1-YFP recombinant protein and the vacuole marker *pUBQ10::RFP-VAMP711* in a XPP cell in a WT plant after a treatment of 90 minutes with endosidine2. **(f)** Cartoon representing GNOM-dependent AUX1 trafficking in WT and *204* mutant plants. White arrowheads: sites of AUX1-YFP recombinant proteins accumulation. Scale bars: 25 μm.

Herex, we show that the stability of GNOM in XPP cells depends on its C-terminal HDS3 domain. The presence of GNOM in this cell type ensures proper recycling of the auxin transporter AUX1, a prerequisite for FC specification and subsequent lateral root formation.

## 4. DISCUSION

GNOM belongs to the family of ADP-ribosylation factors (ARF) guanine nucleotide exchange factors (ARF GEFs), a well-conserve family of proteins among eukaryotes, which is defined by the presence of the SEC7 domain (44, 45). This domain has a nucleotide exchange activity and is responsible for increasing the GTP-bound active state of ARFs, small GTPases associated with membranes to form vesicles (46). There are several subfamilies of ARF GEFs depending on their domain composition (26). These domains can modulate the activity of the SEC7 domain, for instance, favoring homodimerization or interaction with additional partners (47, 48). In concrete, GNOM is part of the Golgi BFA-resistant factor 1 (GBF1) subfamily, which is sensitive to BFA and has three consecutive HDS domains located at its C-terminal half (41, 49). GNOM alleles mutated at the N-terminal half, where the SEC7 domain is located, are embryo lethal (20), but mutant alleles on its C-terminal half can growth adult plant (27, 31). For this reason, we hypothesized whether the C-terminal domain of GNOM, composed of the domains HDS1-3, could have specialized in post-embryonic functions. In this work, we demonstrate that the C-terminal domain of GNOM HDS3 is necessary to stabilize GNOM in cells of the XXP, assuring AUX1 recycling to the membrane. Previous studies, confirmed by our observations, demonstrate that GNOM does not regulate AUX1 trafficking outside the XPP cells (20, 40), indicating that GNOM acquires this function at the XPP cells by the presence of the HDS3 domain. It has been shown that ARFs GEFs can modify the activity of the SEC7 domain by allosteric autoinhibition, favored by the interaction of their structural domains with lipid species located in the membranes (48). In plants, GNOM and the phospholipid flippase ATPase3 have been found to interact with each other to regulate GNOM-dependent PIN1 trafficking (39). This raises the question of whether the HDS3 domain could be involved in the recognition of lipid species present in XPP cells that confer stability of the GNOM protein to regulate the trafficking of AUX1 in this cell type. In addition, we found that part of the HSD3 domain resembles the MONC domain by homology, a conserved domain in Mon2 proteins, proteins with described functions during endomembrane trafficking (50). Sequence similarities between the Mon2 protein in yeast and the non-catalytic regions of GBF1 have been previously described (51). The yeast protein Mon2p regulates trafficking to the vacuole, precisely the pathway affected in mutant 204 (50). Furthermore, Mon2p interacts with amino phospholipid translocases (50), supporting the putative role of HDS3 domains interacting with lipid species. All these findings strongly suggest the involvement of HDS3 during the recognition of lipid species in membranes where GNOM may be regulating vesicle formation. The specific location of these lipid species could determine the areas of activity of the SEC7 domain. Changes in the lipid composition of the endomembrane system of XPP cells could be the key to the differential activity and/or stability of GNOM within these cells led by the presence of the HDS3 domain.

## 5. FIGURE LEYENDS

**FigS1.**
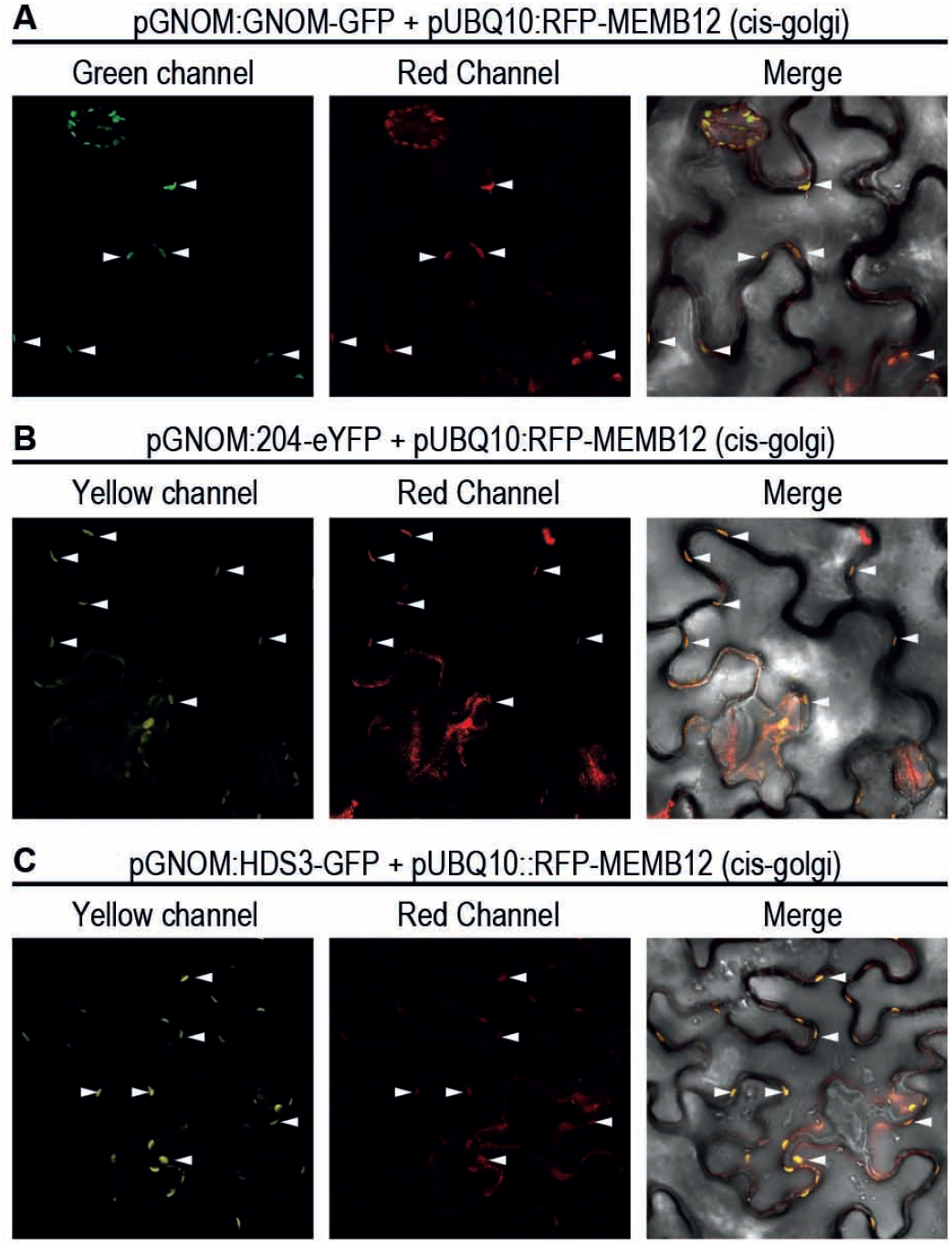
The complete GNOM protein, the truncated 204 version of the GNOM protein and the HDS3 domain of the GNOM protein localize in Golgi aggregates in tobacco leaves. **(a, b, c)** Confocal images showing the accumulation of the GNOM-GFP (a), GNOM-204-eYFP (b) and HDS3-eYFP (c) recombinant proteins and the cis-Golgi apparatus marker *pUBQ10::RFP-MEMB12* in epidermal cells in tobacco leaves. White arrowheads: colocalization points.

**FigS2.**
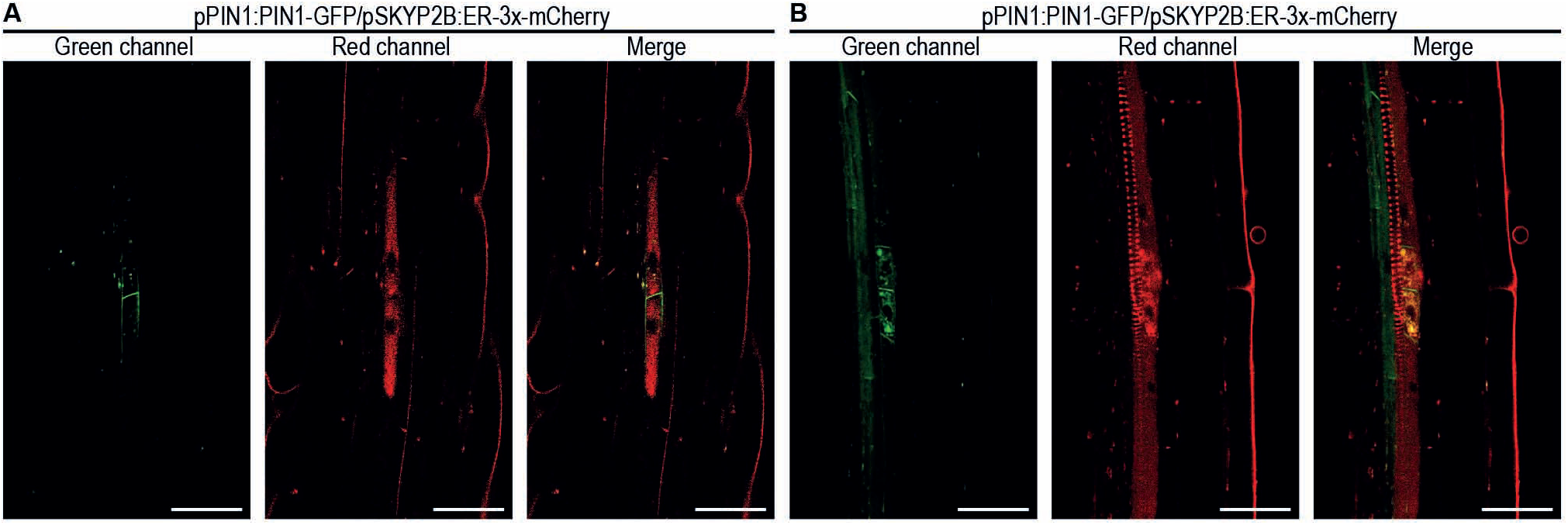
PIN1 is express after founder cell specification. **(a, b)** Confocal images showing the accumulation of PIN1-GFP recombinant protein and the FC marker *pKYP2B::ER-3x-Cherry* in two adjacent FC (a) and after the first asymmetric division (b). Yellow arrowheads: sites of LAX3-YFP recombinant protein accumulation. Scale bars: 25 μm

**FigS3.**
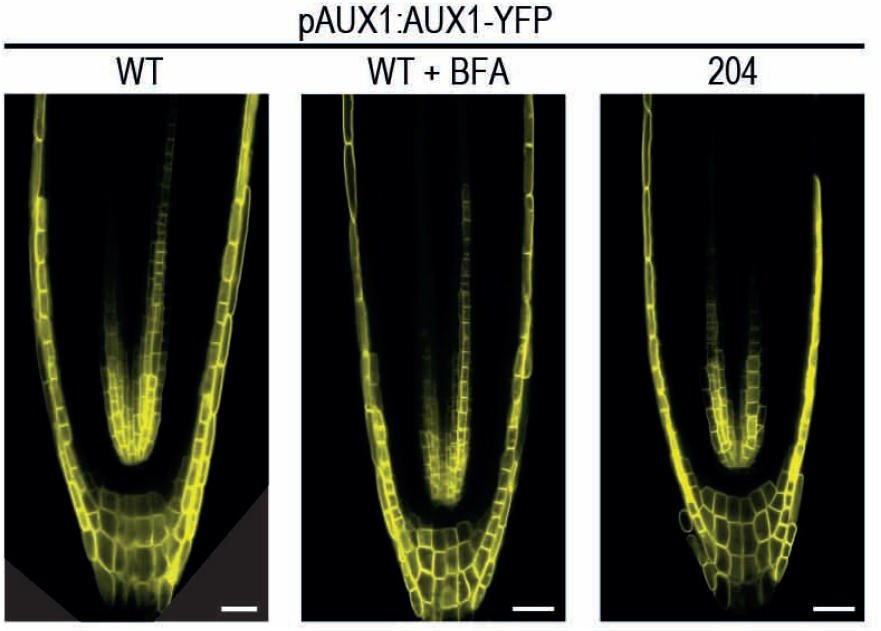
AUX1 remains un-altered in the meristem of the *204* mutant. Confocal images showing accumulation of AUX1-YFP recombinant protein in WT, WT supplemented with BFA and *204* meristems. Scale bars: 25 μm

